# Scarless repair of acute and chronic kidney injury in African Spiny mice (*Acomys cahirinus*)

**DOI:** 10.1101/315069

**Authors:** Daryl M. Okamura, Chris M. Brewer, Paul Wakenight, Nadia Bahrami, Kristina Bernardi, Amy Tran, Jill Olson, Xiaogang Shi, Adrian M. Piliponsky, Branden R. Nelson, David R. Beier, Kathleen J. Millen, Mark W. Majesky

## Abstract

Solid organ fibrosis is a major burden on global health and medical care costs. Muroid rodents of the genus *Acomys* (African Spiny mice) are terrestrial mammals that evolved remarkable abilities to regenerate severe skin wounds without scar formation. However, whether scar-free wound repair in *Acomys* extends beyond skin to vital internal organs is not known. Here, we used two aggressive kidney injury models known to produce severe renal fibrosis and show that despite equivalent acute kidney injury, there was rapid restoration of nephron structure and function without fibrosis in *Acomys* compared to extensive fibrosis leading to renal failure in *Mus musculus*. These results suggest *Acomys* species have evolved genomic adaptations for wound healing that activate regenerative repair pathways not only in skin, but also in vital internal organs. Our findings have important implications for discovering a long-sought evolutionary solution to internal organ injury and regeneration.

## Introduction

Solid organ fibrosis is the result of chronic inflammatory processes and dysregulated wound healing that leads to progressive loss of tissue function and eventual organ failure.^1^ The global health care burden for cumulative loss of vital organ function due to progressive fibrosis is enormous.^2^ There are currently very few treatment options for patients with end stage renal disease or similar degenerative fibrotic conditions in the heart, lung, liver, or other critical internal organs.^3^ Looking to nature for a possible solution, it was reported that adult rodents of the genus *Acomys* (African spiny mice) can shed their dorsal skin as a deterrent to avoid predators and fully regenerate the lost tissue without fibrosis or tissue overgrowth.^4^ The restored skin is complete with hair follicles, sebaceous glands, cartilage, adipose tissue, nerves, and blood vessels in the correct architectural organization.^5,6^ While this remarkable wound healing response in the skin has been examined in some detail, the important question of whether or not scarless regenerative wound repair in *Acomys* species extends beyond skin to vital internal organs remains unanswered.

In the experiments reported here, our objective was to produce injuries to *Acomys* kidney that are known to promote severe fibrotic responses leading to organ failure in murine kidney. Our goal was to determine whether or not scarless, regenerative wound healing observed in *Acomys* skin extends to critical internal organs. We now provide the first reported evidence that scarless wound repair first observed in the skin does indeed extend to a major internal organ in the African spiny mouse. We demonstrate that in two aggressive models of kidney disease, unilateral ureteral obstruction (UUO) and ischemia reperfusion injury (IRI), there was a near complete absence of fibrosis and a rapid regeneration of nephron function in *A. cahirinus.* By contrast, paired groups of *M. musculus* (outbred CD-1 or inbred C57BL/6J) developed severe kidney fibrosis that rapidly progressed to complete renal failure. These studies represent the first step in an evolutionary approach to understand how mammalian wound healing can be uncoupled from the fibrotic response to injury and redirected toward regeneration of complex organ function in mammals.

## Results

### *A. cahirinus* fails to develop fibrosis after UUO injury

Tubulointerstitial fibrosis is the final common pathway of many forms of kidney disease.^1,3,7^ Unilateral ureteral obstruction (UUO) is a reliable and aggressive model of chronic kidney injury and robust interstitial fibrosis. In previously reported studies where the contralateral kidney was removed after 7d of obstruction in *M. musculus,* UUO kidneys were found to have about 50% function. After 14d they become nonfunctional resulting in 100% mortality from kidney failure.^8^ We performed UUO surgeries on *A. cahirinus* and *M. musculus* (outbred CD-1 and inbred C57BL/6J (B6) strains were used) and retrieved injured kidneys (UUO) and contralateral kidneys (NK) at the times indicated in **Fig 1**. We found that even after 14d of obstruction with obvious signs of hydronephrosis (**Fig 1A,B**), the gross anatomic structure and parenchymal thickness (between arrows, **Fig 1C**) were remarkably preserved in *A. cahirinus* compared to *M. musculus* (B6). This preservation of tissue structure was confirmed by the maintenance of relatively normal kidney weights (UUO/NK) in obstructed *A. cahirinus* kidneys compared to rapid declines in kidney weight in *M. musculus* as a a result of progressive fibrosis (*ratio of slopes: m_Mus_/m_Acomys_=-7.5; p=0.03,* **Fig 1C**). There were no significant differences in uninjured contralateral kidney weights between *A. cahirinus* and *M. musculus* (data not shown).

**Figure 1.**
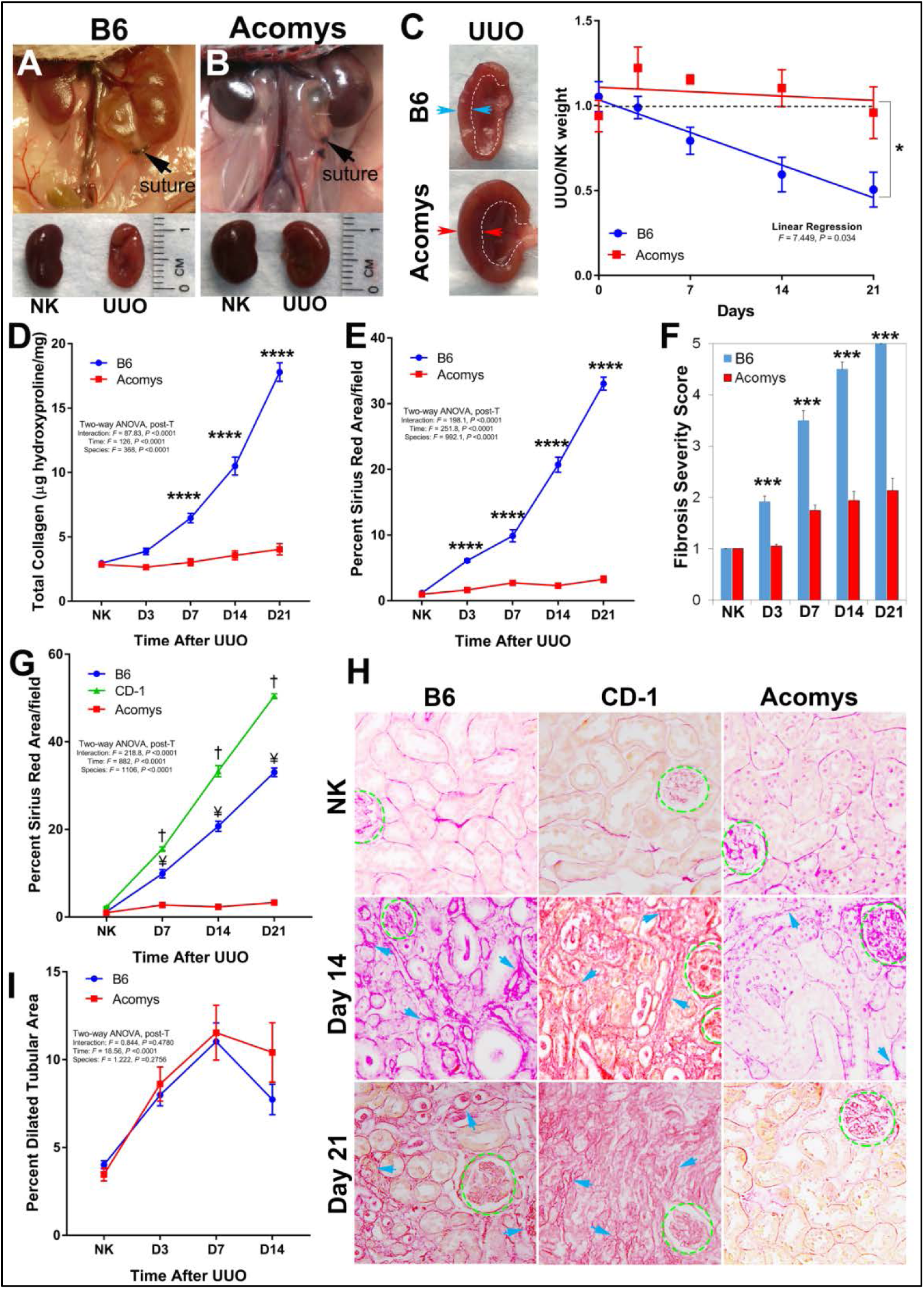
Absence of fibrosis after severe obstructive injury in *A. cahirinus*. **(A,B)** (Upper) The ureter from the left kidney was ligated (arrow) to produce obstructive injury in both *A. cahirinus* and C57BL/6J (B6) mice. (Lower) Upon gross inspection, contralateral kidneys (NK) from the two species are similar in length but *A. cahirinus* UUO kidneys appear less damaged than B6 UUO kidneys after 14d of obstruction; **(C)** Left panel demonstrates preservation of parenchymal thickness (greater distance between arrows) in *A. cahirinus* kidney with renal pelvis noted by dotted white line. The graph demonstrates the best fit line of the ratio of UUO:NK kidneys for each time point and the slopes were analyzed by linear regression (n=6- 10/time point); **(D)** Total collagen content was measured by micrograms hydroxyproline per mg wet kidney weight. Graph summarizes total collagen measurement for B6 and *A. cahirinus* NK and UUO kidneys, (n=6-8/time point for each group); **(E)** Graph summarizes image analysis of picrosirius red staining for each UUO time point (n=6/time point for each group); **(F)** Masson trichrome sections were blindly scored for IFTA and inflammation (*see Methods*, n=6/time point for each group); Graph summarizes IFTA scores (fibrosis severity score) for each time point; **(G)** UUO was performed on outbred CD-1 mice (green)(n=3-4/time point) and the development of fibrosis was compared to B6 (blue) and *A. cahirinus* (red). Graph summarizes image analysis of picrosirius red staining with **(H)** representative digital images (400x). Glomeruli are outlined (dotted green). Arrows demonstrate Sirius red staining of interstitial matrix. **(I)** Dilated tubular area was measured in Masson trichrome sections; Graph summarizes image analysis of tubular dilation area in B6 and *A. cahirinus* after UUO (n=6-7/time point for each group): *B6 vs A. cahirinus*: * p<0.05, **p<0.01, ***p<0.001, ****p<0.0001; *CD-1 vs A. cahirinus*: † p<0.0001; *B6 vs CD-1*: ¥ p<0.0001).

Progression of fibrosis was monitored by three different assays. Total collagen levels were measured as hydroxyproline content per wet kidney weight. Kidney collagen levels increased rapidly in *M. musculus* while *A. cahirinus* exhibited no significant change from the uninjured contralateral kidney (NK) (n=6-8, **Fig 1D**). Remarkably, even out to 21d of obstruction, there were no significant differences in total collagen levels between UUO kidney and uninjured contralateral kidney (NK) in *A. cahirinus* (**Fig 1D**; *Acomys:NK vs D3-21, NS*). Computer-assisted image analysis of picrosirius red-stained kidney tissue sections demonstrated a nearly complete absence of interstitital matrix fibrosis at each time point after UUO injury in *A. cahirinus* even out to 21d of obstruction (*Acomys*: *NK vs D3-21, NS*), compared to extensive fibrosis in *M. musculus* kidneys (**Fig 1E**, **Supp Fig 1**). Interstitial fibrosis, inflammation, and tubular atrophy (IFTA) were blindly scored on Masson’s trichrome stained sections. We found that IFTA scores were markedly reduced in *A.cahirinus* compared to *M. musculus* (B6) despite chronic obstructive injury in both species (**Fig 1F**, **Supp Fig 2**). In order to test our findings in an outbred strain of *M. musculus*, we performed UUO surgeries on CD-1 mice and measured fibrosis severity by picrosirius red staining. Of interest, we found even more dramatic increases in interstitial fibrosis in CD-1 mice producing even greater differences in fibrotic tissue areas when compared to *A. cahirinus* (**Fig G,H**; *p<0.0001*). All together, these results demonstrate that, in contrast to either inbred or outbred *M. musculus* strains, *A. cahirinus* does not develop fibrotic tissue in response to severe chronic obstructive kidney injury.

### *A.cahirinus* maintains tubular integrity and modifies myofibroblast accumulation after UUO injury

In order to quantify the degree of obstructive tubular injury, we measured the dilated tubular area in *A. cahirinus* and *M. musculus* (B6) at 3d through 14d after UUO. As expected from the obvious hydronephrosis seen in **Fig 1A,B**, we found that tubular dilation increased significantly in both species following UUO compared to the contralateral uninjured kidney, and peaked at 7d (**Fig 1I**; *p<0.05*). Importantly, there were no significant differences in the extent of tubular dilation between *A. cahirinus* and *M. musculus* at any of the time points examined (**Fig 1I**). Activated myofibroblasts (positive for smooth muscle α-actin, *Acta2*) are a significant source of collagen-rich extracellular matrix produced during kidney fibrosis. Chronic tubular injury is known to promote the production of intrarenal profibrotic cytokines that activate myofibroblasts.^9^ While *Acta2* immunolabeling increased after UUO in both species, *M. musculus* exhibited higher levels of *Acta2*+ myofibroblasts compared to *A. cahirinus* (**Fig 2A,B**; *p<0.01*). In contrast to the lack of significant fibrosis in *A. cahirinus* after UUO (**Fig 1D,E**), there was a significant increase in *Acta2+* myofibroblasts at D14 after UUO compared to both NK and D3 time points (**Fig 2B**, *p<0.05*). These results suggest that the absence of interstitial matrix deposition in *A. cahirinus* after UUO injury is not due to the absence of myofibroblasts.

**Figure 2.**
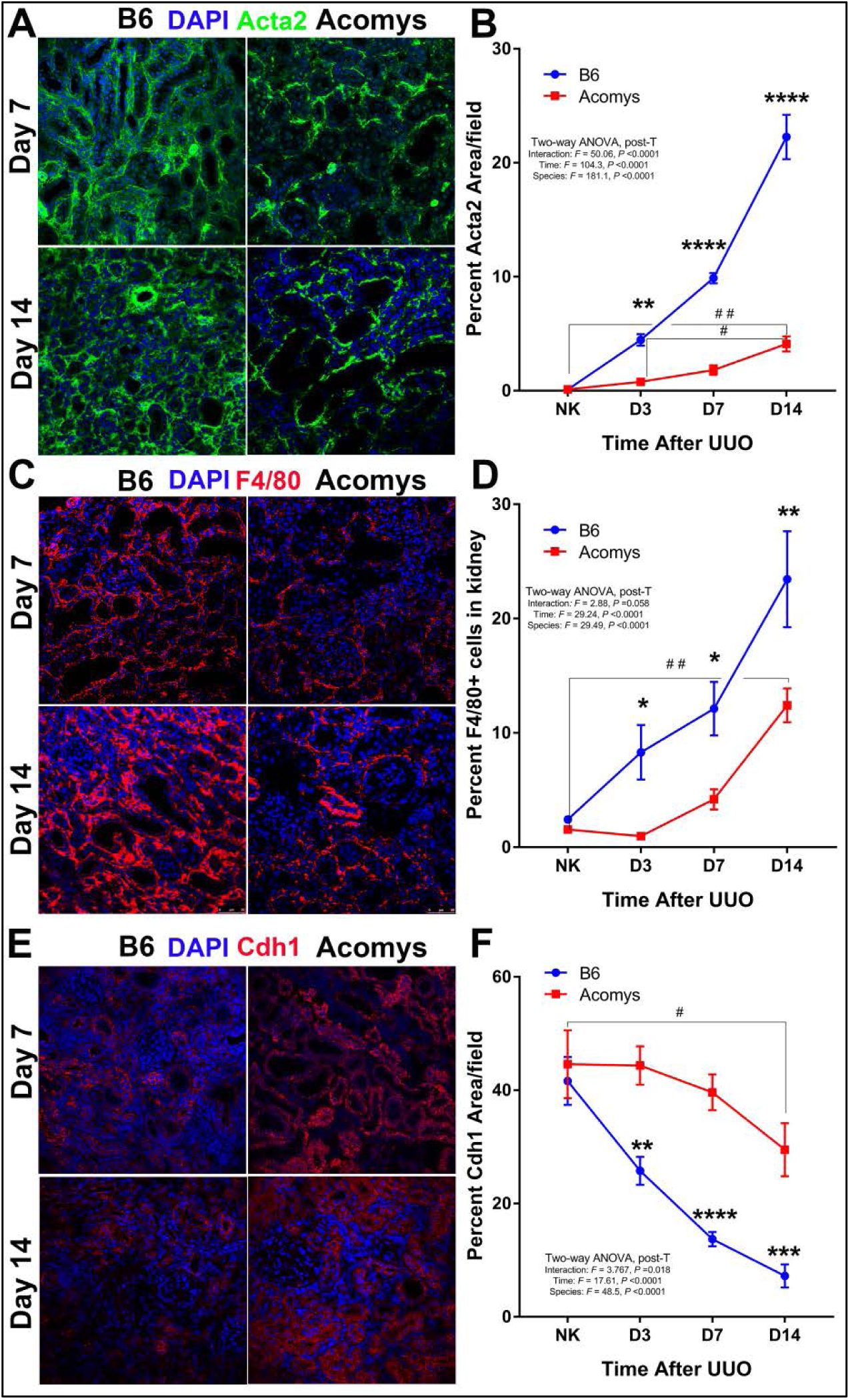
Myofibroblast formation and macrophage infiltration do not generate a fibrotic response in *A. cahirinus*. **(A)** Smooth muscle alpha actin (*Acta2*) expression was investigated by immune-confocal microscopy at days 3, 7, and 14 after UUO. Representative digital images (400x) *Acta2* expression (green) for B6 and *A. cahirinus* at days 7 and 14 after UUO; **(B)** Graph summarizes image analysis for *Acta2* at each time point (n=6/time point for each group). **(C)** F4/80 macrophage infiltration was examined by confocal microscopy and quantified by flow cytometry. Representative digital images (400x) of F4/80 (red) expression at days 7 and 14 after UUO. **(D)** Flow cytometry was performed on single cell suspensions from whole kidney and analyzed for F4/80 expression. Graph summarizes percent F4/80+ cells at days 3, 7, and 14 after UUO analyzed by FACS (n=3-4/time point per group). **(E)** Tubular integrity was examined by confocal microscopy for *Cdh1* (E cadherin). Representative digital images (400x) of *Cdh1* (red) expression for B6 and *A. cahirinus*. **(F)** Graph summarizes image analysis results for *Cdh1* levels at each time point (n=6/time point for each group). (B6 (blue) vs. *A. cahirinus* (red): *p<0.05, **p<0.01, ***p<0.001, ****p<0.0001; between time points # p<0.05, ## p<0.01)

Chronic inflammation with a predominance of macrophages is a characteristic finding in organ injury and is strongly correlated with tissue fibrosis.^10^ In order to quantify macrophage infiltration, whole kidneys were enzymatically digested into single cell suspensions and analyzed for F4/80 expression by flow cytometry (**Fig 2C,D**). As expected, the number of F4/80 macrophages increased in *M. musculus* UUO kidneys with advancing obstruction (7d and 14d) compared to contralateral normal kidneys (*p<0.001*). In *A. cahirinus,* increases in macrophage content were both delayed and diminished over the same time course (**Fig 2D**). Similar to *Acta2* data from *A. cahirinus* kidneys, there was a significant increase in F4/80+ macrophages at D14 compared to NK and D3 time points (**Fig 2D**, *p<0.001*). In comparing *A. cahirinus* with *M. musculus* kidneys, significant reductions in F4/80+ macrophage content were found at each time point examined (*p<0.05*) but less dramatic than the fibrosis (**Fig 1D,E**) and myofibroblast (**Fig 2B**) data. These results suggest that the unique absence of fibrosis in *A. cahirinus* is not due to a complete absence of a chronic inflammatory response or an absence of myofibroblasts in injured kidney tissues but suggests an evolutionary adaptation in wound repair.

Tubular integrity is strongly correlated with nephron function and can serve as a histological surrogate of whole kidney function.^11,12^ *Cdh1* (E-cadherin) is an indicator of tubular cell integrity and polarity whose expression is lost with ongoing obstructive injury leading to loss of tubular architecture.^13^ As expected for *M. musculus* there was progressive loss of *Cdh1* expression with each time point after UUO compared to the contralateral normal kidney (**Fig 2E,F**). However, in *A. cahirinus*, there were no significant changes in *Cdh1* protein levels with advancing obstructive injury until D14 compared to the contralateral uninjured kidney. There were no differences in *Cdh1* expression levels in uninjured contralateral kidneys between *A. cahirinus* and *M. musculus* (**Fig 2F**). Thus, despite severe chronic obstruction, *Cdh1* protein levels were maintained in *A. cahirinus* while becoming significantly decreased in *M. musculus* injured kidneys with each time point (**Fig 2F**). All together, these results demonstrate that despite equivalent tubular dilation with obstruction (**Fig 1K**), we found significant attenuations in myofibroblast activation, macrophage infiltration, and loss of *Cdh1* in *A. cahirinus* kidneys that were correlated with preservation of tubular integrity and lack of interstitial fibrosis compared to *M. musculus*.

### Renal fibrosis is reduced in *A. cahirinus* despite equivalent ischemic injury

Although the UUO model is useful in the study of renal fibrosis, it is not directly translatable to human kidney disease. By contrast, ischemic kidney injury is one of the most common causes of acute kidney injury in humans. We sought to test our findings in a second model of severe kidney injury with unilateral ischemia-reperfusion injury (uni-IRI) following prolonged 40 min of ischemia. One of the questions arising from our results with the UUO model was whether *A. cahirinus* was able to resist acute kidney damage rather than alter the subsequent fibrogenic cascade. Therefore, we performed uni-IRI with a simultaneous contralateral nephrectomy (uni-IRI+Nx) on *A. cahirinus* and *M. musculus* and sacrificed them at 24h after surgery in order to correlate kidney function with histology after severe acute injury. We found a significant elevation in blood urea nitrogen (BUN) levels 24h after uni-IRI+Nx in both *A. cahirinus* and *M. musculus* (**Fig 3A**). Importantly, these acutely elevated BUN levels were not significantly different between species (**Fig 3A**). In fact, there was a trend towards higher BUN levels in *A. cahirinus* (BUN: *Acomys* vs *Mus*, 129±24 vs 102±16 mg/dL). H&E sections on the Uni-IRI-Nx kidneys from *A. cahirinus* and *M. musculus* at 24h were analyzed for tubular cell necrosis, casts, and dilation and assigned cumulative tubular injury scores. Consistent with our kidney function data, we found no differences in tubular injury scores at 24h between *A. cahirinus* and *M. musculus* (**Fig 3B,C**) and therefore conclude that both species experience equivalent levels of acute ischemic injury and tissue damage after prolonged renal ischemia as assessed both histologically and functionally.

**Figure 3.**
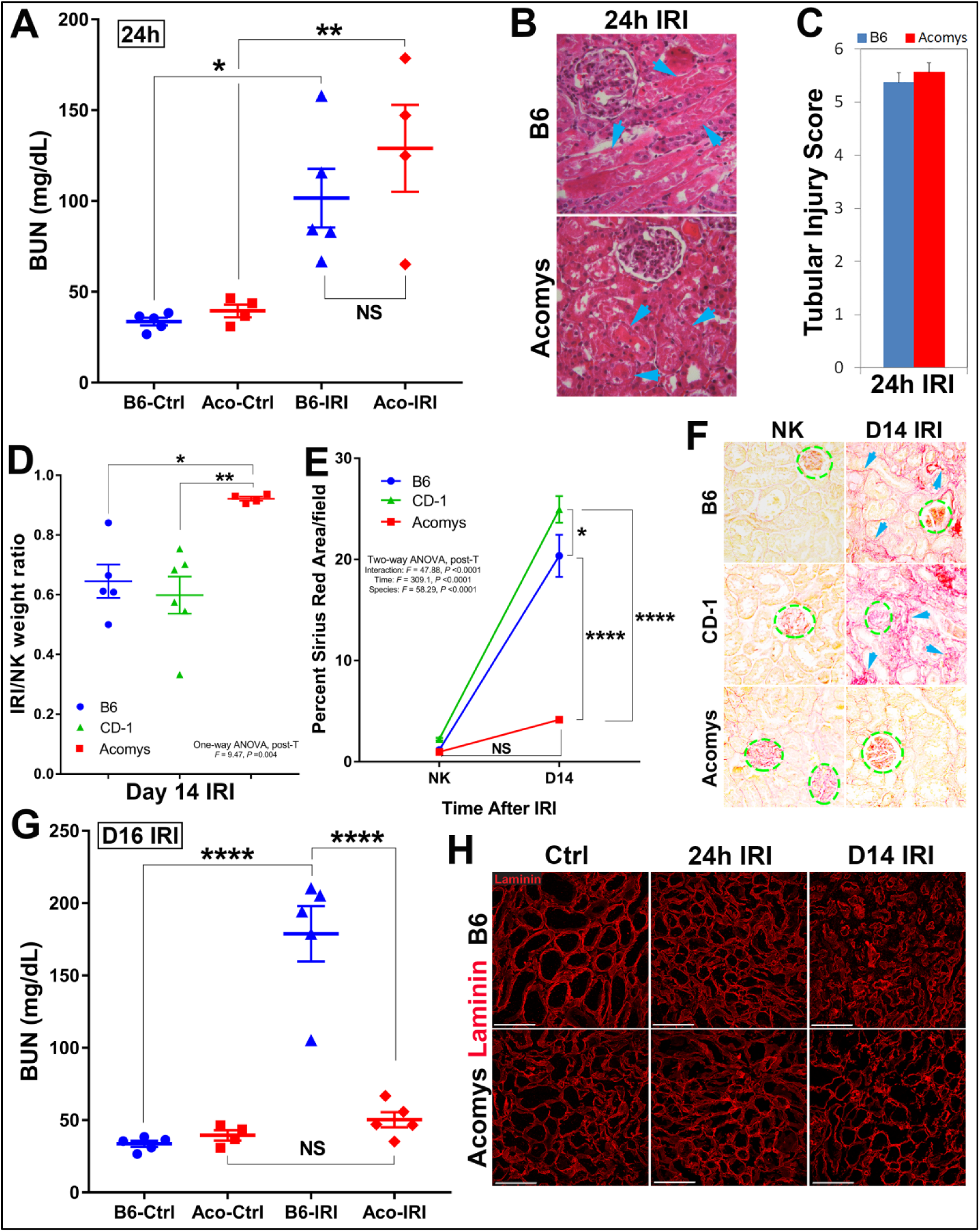
Near complete recovery of nephron function after severe ischemic injury in *A. cahirinus*. **(A)** B6 and *A. cahirinus* underwent unilateral IRI (uni-IRI) with contralateral nephrectomy and sacrificed at 24h. BUN levels were assessed in uninjured animals (Ctrl) and those sacrificed at 24h (n=4-5/time point per group):): NS–not significant, * p<0.05, **p<0.01. Note that both species exhibited equivalent severe acute kidney damage in response to ischemic injury. **(B)** H&E stained sections were blindly scored for tubular injury (*see Methods*, n=4-5/group); Representative H&E digital images (400x) at 24h from IRI kidneys. Arrows indicate tubular casts and necrotic cell debris. **(C)** Graph summarizes tubular injury scores at 24h after IRI. **(D)** In order to determine effect of acute injury on repair, uni-IRI was performed without nephrectomy on B6, CD-1, and *A.cahirinus*. Graph summarizes data on IRI/contralateral (NK) kidney weight ratio at time of sacrifice at 14d (n=4-6/group): * p<0.05, **p<0.01. **(E)** Graph summarizes image analysis of picrosirius red staining for each UUO time point (n=5-6/time point for each group); (B6 (blue), CD-1 (green), *A.cahirinus* (red): * p<0.05, ****p<0.0001. **(F)** Representative picrosirius red digital images (400x). Glomeruli are outlined (dotted green). Arrows demonstrate Sirius red staining of interstitial matrix. **(G)** In order to determine functional recovery, uni-IRI was performed, the contralateral kidney was removed at day 14 and kidney function was monitored until sacrifice at day 16. BUN levels were determined in uninjured animals (Ctrl) and those sacrificed at day 16, 2 days after contralateral nephrectomy (IRI). Note the nearly complete recovery of nephron function in the injured *A. cahirinus* kidneys at 16d compared to the high BUN levels indicative of kidney failure in B6 injured kidneys: NS-not significant, ****p<0.0001. **(H)** Laminin (red) immunostaining of tubular epithelial basement membrane architecture at days 0, 1, and 14 after IRI injury (400x; scale bars 100um). Note *A. cahirinus* kidney at day 14 (D14 IRI) strongly resembles uninjured kidney (Ctrl) in tubular basement membrane architecture, while B6 basement membranes demonstrate collapse and thickening with advancing fibrosis. Scale bars; 100μm.

In order to investigate the ability of *A. cahirinus* to repair acute kidney injury, we performed uni-IRI without contralateral nephrectomy to allow long term survival in both species and then sacrificed animals at 14d to assess kidney structure by histology. Remarkably, despite severe acute ischemic injury, we found a near complete absence of fibrosis by picrosirius red staining, and a robust preservation of renal mass, in *A. cahirinus* compared to *M. musculus* (either outbred CD-1 or inbred B6) (**Fig 3D-F**). For example, IRI/contralateral kidney weight ratios at 14d were maintained in *A. cahirinus* (0.92±0.02) compared to almost 40% loss of renal parenchymal mass to fibrosis in *M. musculus* (**Fig 3D**). Likewise, fibrosis severity measured by picrosirius red staining indicated a near total absence of fibrosis in *A. cahirinus* compared to either CD-1 or B6 strains of *M. musculus* (**Fig 3E,F**).

To assess the degree of functional restoration in the uni-IRI damaged kidney, we performed uni-IRI followed by a contralateral nephrectomy at day 14 and then measured kidney function over the next 2 days. Importantly, we found striking and reproducible differences in 16d BUN levels between *A. cahirinus* and *M. musculus* (**Fig 3G**). Consistent with a near complete absence of tubular damage and interstitial fibrosis at 14d, we found that BUN levels were nearly normal in *A. cahirinus* indicative of almost complete restoration of kidney function by 16d after IRI (compare **Fig 3A** to **Fig 3G**). By contrast, *M. musculus* BUN levels were substantially increased indicative of progressive renal failure (**Fig 3G**). Likewise, staining for the tubular epithelial cell basement membrane protein laminin showed progressive disruption and thickening by 14d in *M. musculus* kidney (B6) consistent with tubular atrophy while basement membrane structures at 14d in *A. cahirinus* kidney strongly resembled normal uninjured kidney (Ctrl) (**Fig 3H**) consistent with our Sirius red results (**Fig 3E**). The removal of necrotic and cellular debris is an important precursor in wound repair and tissue regeneration.^2^ Intraluminal casts and debris were quantitated from kidney sections and showed that both species exhibited abundant casts/debris at equivalent levels by 24h after IRI (**Fig 4A,B,C**, arrows). At later time points, *M. musculus* retained these intraluminal debris while *A. cahirinus* efficiently cleared them from the tubular network (**Fig 4A, D-G**). Histologic examination by periodic acid Schiff (PAS) stains demonstrated that there were similar levels of tubular necrosis and tubular casts seen in the corticomedullary junction at 24h after severe IRI in both *A. cahirinus* and *M. musculus* (**Supp Fig 3A-C**). However, what was strikingly different in *A. cahirinus* was the abundance of polymorphonuclear cells and other nucleated cells within intraluminal tubular casts (**Supp Fig 3D,E**, arrows) that was seen much less frequently in *M. musculus* (**Supp Fig 3B**, arrow). By 72h after IRI, tubular casts, dilation, and interstitial inflammation progressed in *M. musculus* (**Supp Fig 3F,G**) while in *A. cahirinus* the intraluminal cellular debris had been removed and replaced by highly nuclear, somewhat disorganized tubular structures (**Supp Fig 3H**) with flattened epithelial cells suggesting progression toward a more dedifferentiated state (**Supp Fig 3I,J**, arrows). By 7d, monocytic infiltrates, tubular damage and interstitial fibrosis continue to progress in *M. musculus* compared to the appearance of defined tubular structures with open lumens in *A. cahirinus* (**Supp Fig 3M,N**) and PAS-positive brush border structures signifying mature differentiated tubular epithelial cells (**Supp Fig 3O**). These findings confirm that the response to severe acute kidney injury in *A. cahirinus* does not lead to the progressive, degenerative fibrotic response characteristic of *M. musculus* and human kidneys, but instead results in near complete restoration of nephron structure and function.

**Figure 4.**
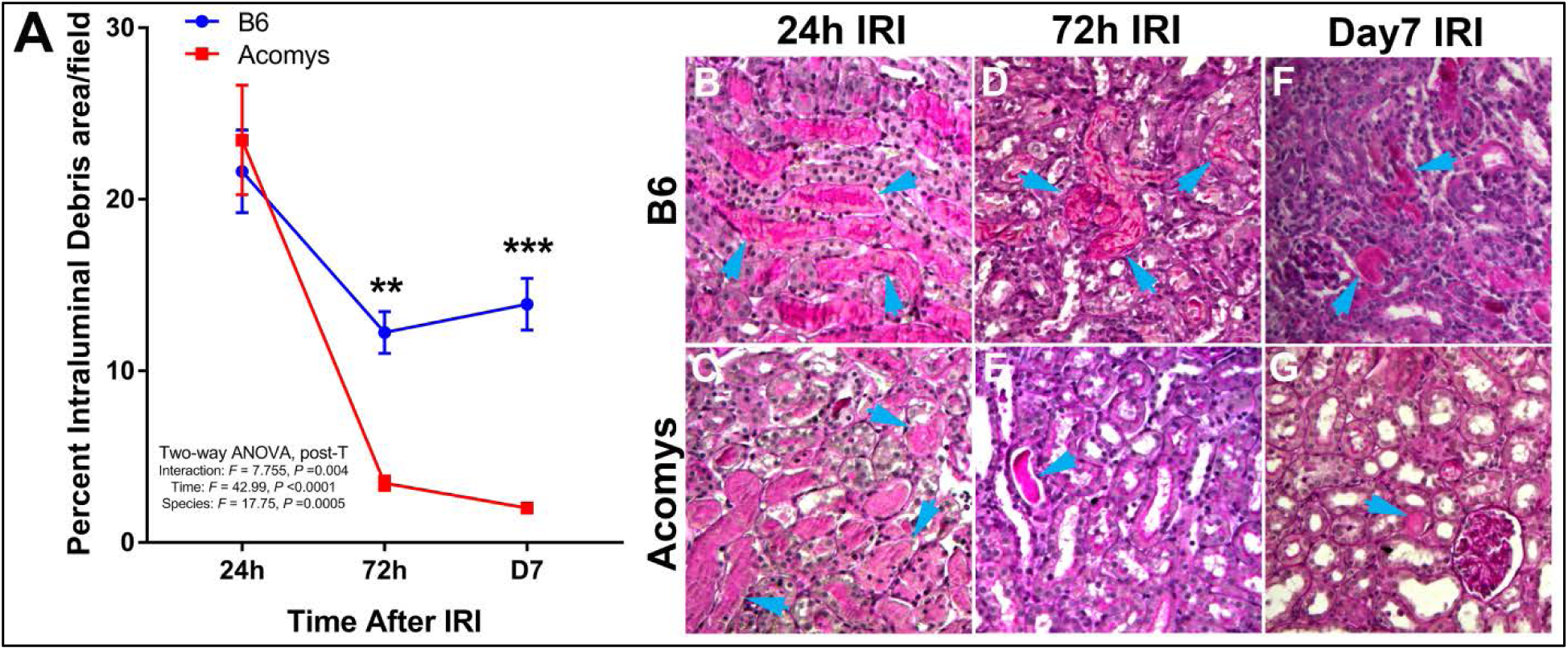
Rapid clearance of tubular debris after severe ischemic injury in *A. cahirinus*. Tubular casts and debris were identified on Periodic acid Schiff (PAS) stain in unilateral IRI kidneys at 24h, 72h, and 7 days in B6 and *A. cahirinus*. **(A)** Graph summarizes image analysis results for tubular casts and intraluminal debris after IRI. (B6-blue, *A. cahirinus-* red; *B6 vs A. cahirinus*: **p<0.01, ***p<0.001). Representative fields from 200x images demonstrate representative tubular casts and intraluminal debris (arrow) in IRI kidneys at 24h **(B,C)**, 72h **(D,E)**, and 7 days **(F,G)** in B6 and *A. cahirinus* kidneys. Arrows demonstrate areas of intraluminal debris/casts.

## Discussion

Using two different and highly aggressive forms of experimental kidney injury, we show that there was a near complete absence of interstitial renal fibrosis and a remarkable restoration of kidney function in *A. cahirinus* compared to either inbred (B6) or outbred (CD-1) strains of *M. musculus*. These remarkable differences in wound healing responses were not due to the failure of our injury models to produce acute tissue damage in *A. cahirinus* kidneys since histological and functional assays showed equivalent tissue injuries in the first 24-72h after UUO or IRI surgeries in both species. Particularly striking was the almost complete restoration of normal kidney function, assessed by blood urea nitrogen levels, by 14d after severe ischemia-reperfusion injury in *A. cahirinus* (**Fig 3G**). By contrast, in parallel experiments with *M. musculus,* injured kidneys were severely fibrotic and progressing rapidly towards complete kidney failure (**Fig 3G**).

Muroid rodents of the genus *Acomys* (Spiny mice) have evolved the ability to shed their dorsal skin to avoid predation and then to completely regenerate the lost skin tissues without fibrosis or scar formation (4-6). Mechanical and histological assays showed that *Acomys* skin is specialized to be structurally fragile and prone to tear under low tensile forces.^4^ Therefore it cannot be assumed that the regenerative response to tissue injury in the skin, the first target of predatory attacks in the wild, necessarily extends to internal organs in *Acomys* species. A similar ear skin regenerative response was previously reported for the MRL/MpJ strain of mice.^14^ However, multiple attempts to determine if regenerative wound healing extended to internal organs, including kidneys, of these mice were generally negative.^15-17^ Thus, our results on the striking absence of fibrotic tissue formation in *A. cahirinus* kidney suggest that the regenerative wound healing response previously described in the skin^4-6^ may indeed be a systemic property that extends to critical internal organs in this species.

Acute kidney injury (AKI) initiates a fibrogenic cascade that leaves patients at high risk for developing chronic kidney disease and progressive loss of renal function.^18-20^ Although elegant studies in *M. musculus* have produced substantial insights into the pathogenesis of renal fibrosis, translating these findings into therapeutic solutions has been poor. Wound healing in most adult mammals, including humans, is a process of repair that ultimately replaces functional tissue with a collagen-rich extracellular matrix resulting in a corresponding loss of organ function. By contrast, some fish and amphibian species can fully regenerate tissue damage and restore organ function after amputation or severe tissue injuries.^21^ In the zebrafish kidney, for example, there is evidence of formation of new nephrons after gentamicin nephrotoxicity.^22^ However, in adult mammals there are no reports of nephron formation *de novo* after kidney injury. We now provide evidence for a potentially transformative new mammalian model for kidney disease that has evolved a distinctly different wound healing response to kidney injury than the currently studied mouse, rat, or human, models. If confirmed for other organs, an in-depth analysis of the molecular basis for scar-free wound healing in Acomys species could be a gateway for novel anti-fibrotic therapies.

While our results demonstrating nearly complete restoration of kidney function after severe IRI injury (**Fig 3G**) are consistent with the likelihood there is nephron regeneration in *A. cahirinus,* they are not currently sufficient to prove this. In models of true vertebrate regeneration, such as axolotl limb,^23^ zebrafish heart,^24^ or *Acomys* skin,^5^ tissue mass is removed by surgical resection and the lost tissue is fully restored both structurally and functionally. Although tissue damage produced by 40 minutes of renal ischemia/reperfusion is extensive (**Fig 3A-C**, **Fig 4**) and restoration of kidney function is remarkably robust (**Fig 3G**), further work is required to definitively establish whether new nephrons are being formed in this model. The most robust phenotypic difference between *A. cahirinus* and *M. musculus* in our study is the near complete absence of fibrotic tissue formed in injured *A. cahirinus* kidneys after either UUO or IRI procedures. The extent to which lack of fibrosis is sufficient to explain the differences in wound healing outcomes reported here, or whether other changes in the *A. cahirinus* genome are essential for regenerative wound repair, are important questions for future work.

## Acknowledgments

This work was supported by a grant from the W.M. Keck Foundation, the US National Institutes of Health grant 1R21OD-023838, the Loie Power Robinson Stem Cell & Regenerative Medicine Fund, and the Seattle Children’s Research Institute.

## Author Contributions

DMO, CMB, PW, NB, KB, AT, XS, JO, AMP, BRN performed the experiments. DMO, DRB, MWM designed the experiments. DMO, CMB, MWM wrote the manuscript. DMO, KJM, MWM obtained funding for the project. All authors made critical input into editing the manuscript.

## Author Information

Correspondence and requests for materials should be addressed to one of the following: DO; KJM; or MWM.

## Supplemental Figure Legends

**Supplemental Figure 1.**
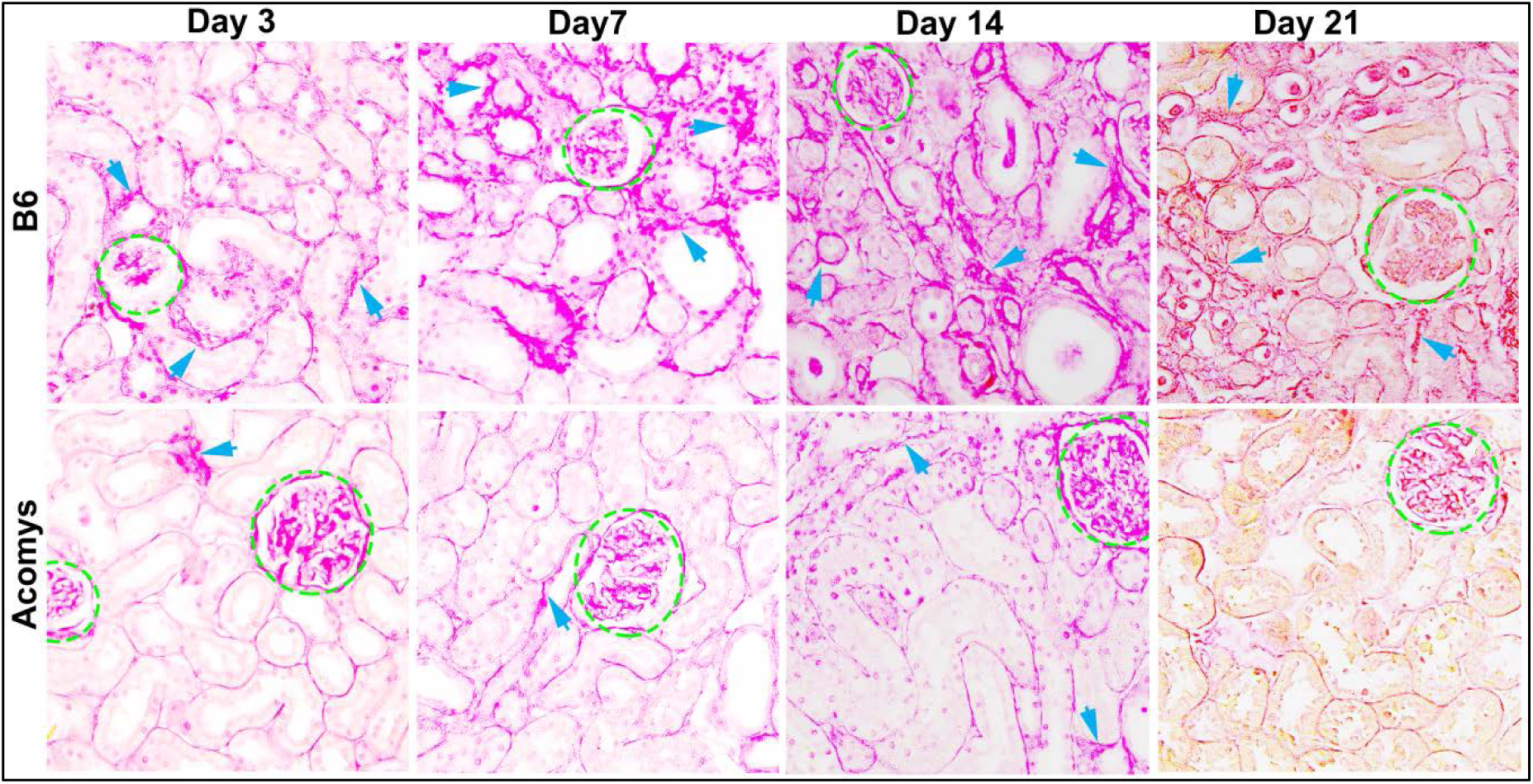
Picrosirius red staining in B6 and *A. cahirinus*. Representative picrosirius red digital images (400x) in B6 and *A. cahirinus* after UUO. Glomeruli are outlined (dotted green). Arrows demonstrate Sirius red staining of interstitial matrix.

**Supplemental Figure 2.**
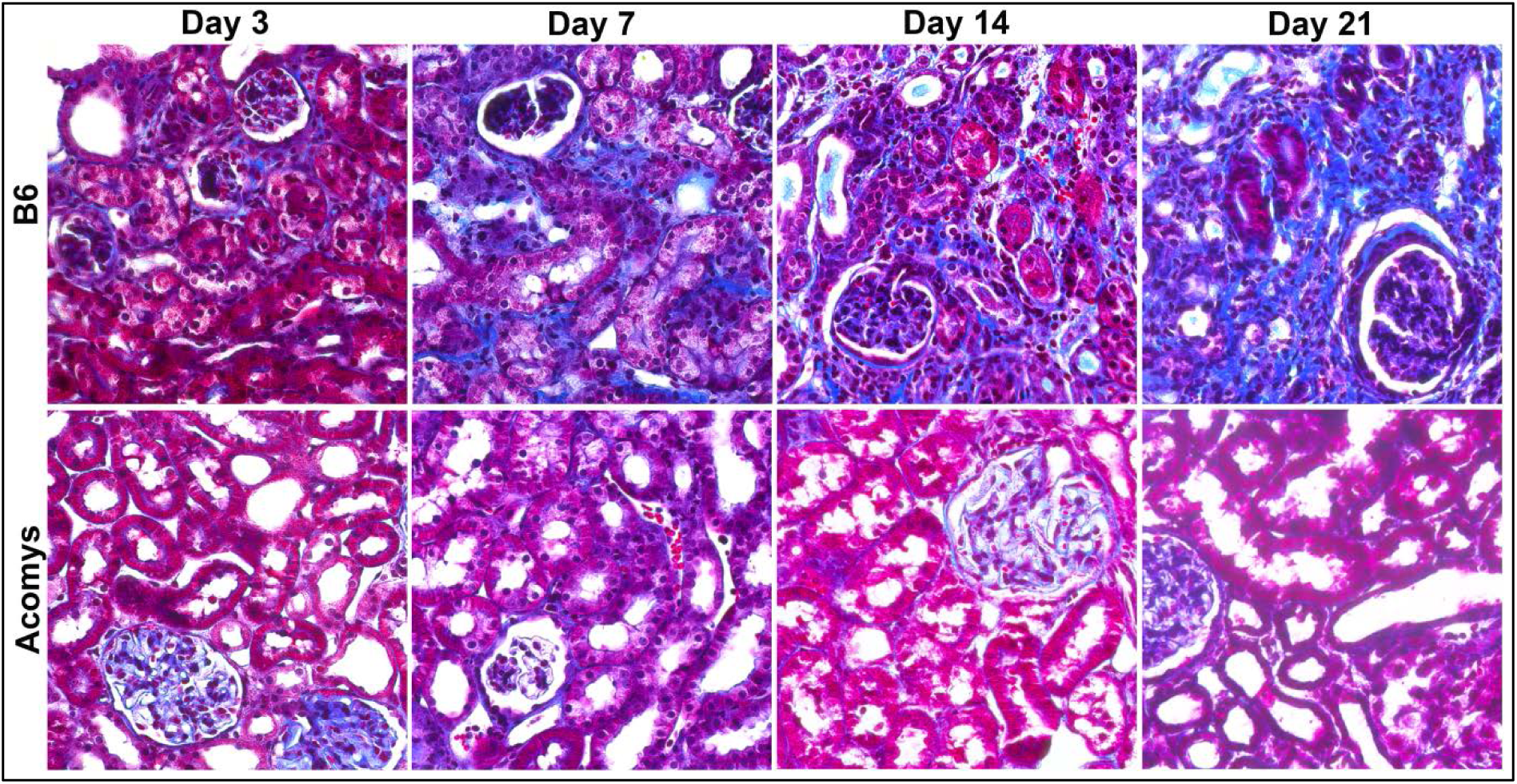
Fibrosis severity score in B6 and *A. cahirinus*. Masson trichrome sections from were blindly scored for IFTA and inflammation (*see Methods*, n=6/time point for each group). Representative Masson trichrome digital images (400x) after UUO.

**Supplemental Figure 3.**
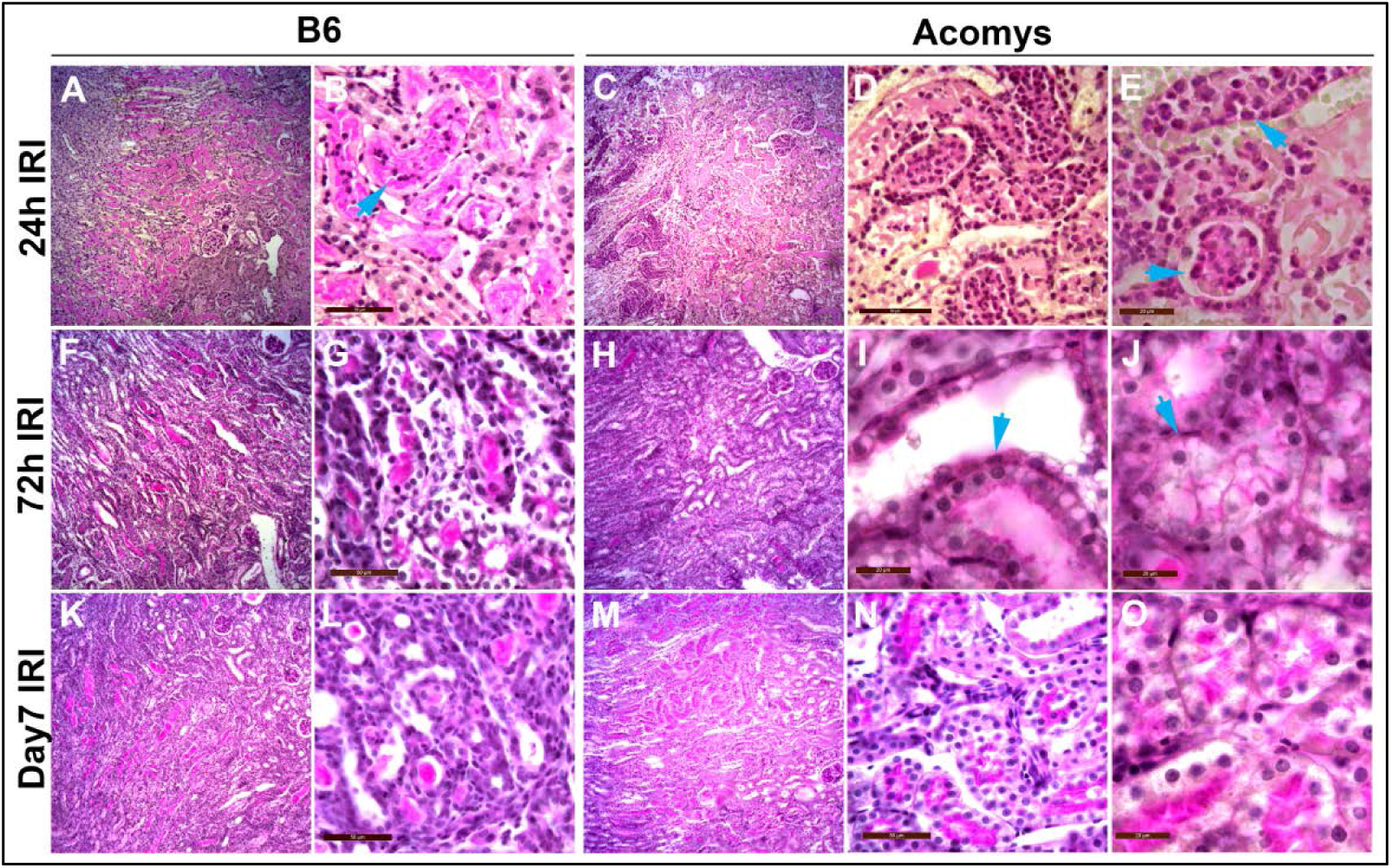
Restoration of nephron structure after severe ischemic injury in *A. cahirinus*. Periodic acid Schiff (PAS) stain was performed on unilateral IRI kidneys at **(A-E)** 24h, **(F-J)** 72h, and **(K-O)** 7 days in B6 and *A. cahirinus*. Low power images (100x) in B6 demonstrate progression of **(A)** necrotic kidney injury to **(F,K)** tubular casts and loss of tubular structure with interstitial inflammation. In comparison, low power images in *A. cahirinus* show similar necrotic injury **(C),** with accelerated repair and restoration of nephron structure **(H,M)**. At 24h, polymorphonuclear cells can be seen in tubular lumens and casts in B6 (arrow **B**), and to an even greater extent in *A. cahirinus* (arrows **E**). At 72h, tubular casts are nearly resolved (arrow, I), and flattened, dedifferentiated tubular epithelial cells **(I)** and tubular lumens in repair (arrow **J**) can be seen in *A. cahirinus*. At 7 days, tubular casts and interstitial inflammation remain in B6 **(L)**, compared to PAS positive brush border cells in *A. cahirinus* **(N,O)**.

## Concise Methods (to be in online supplement)

### Experimental Design

We utilized two models of kidney injury to investigate the differences in wound healing and fibrosis: Unilateral ureteral obstruction (UUO) and ischemia reperfusion injury (IRI). C57BL/6J (B6) and CD-1 mice were used as inbred and outbred strains of *M. musculus,* respectively. Surgery was performed on male animals between 3 and 6 months of age. UUO surgeries were performed as previously described^25,26^ (n =6-8/group), and animals were sacrificed at 3, 7, 14, and 21 days after surgery. Mice received isoflurane anesthesia (5% induction; 1-3% maintenance) in oxygen through a precision vaporizer (Portable Anesthesia Machine, PAM; Molecular Imaging Products, Bend, OR, USA).

Unilateral IRI (uni-IRI) surgeries were performed as previously described^25,26^ except that the vascular pedicle was clamped for 40 minutes (n=5-6/group), and animals were sacrificed at 24h, 72h, 7 days, and 14 days after surgery. In order to assess initial injury, the contralateral kidney was removed at the time of uni-IRI surgery and sacrificed at 24h after surgery. In order to quantify kidney function in the uni-IRI kidney after initial injury, a contralateral nephrectomy was performed 14 days post-surgery as previously described.^25^ Blood was drawn daily until sacrifice at 2 days post-nephrectomy. All procedures were performed in accordance with the guidelines established by National Research Council Guide for the Care and Use of Laboratory Animals and approval of our Institute Animal Care and Use Committee (IACUC). Contralateral, UUO, and IRI kidneys were harvested and processed for RNA and protein extraction and histological studies as previously described.^25-27^ Frozen tissue samples were stored at-80°C.

### Collagen content

Hydroxyproline content of kidney tissue (μg of hydroxyproline per mg of wet weight kidney section) was measured by acid hydrolysis of the tissue section using procedures established in our laboratory.^25-27^

### Histological examination

Immunohistochemical staining was performed on sections of paraffin-embedded tissue or cryosections of snap-frozen tissue using procedures established in our laboratory with VECTASTAIN *Elite* ABC Kits (Vector Laboratories, Inc.) and AEC Substrate Chromogen K3464 (Dako Corp.). Sections were blocked with Avidin/Biotin blocking kit (Vector Laboratories, Inc.). Computer-assisted image analysis was performed on 6 randomly selected 400x magnified images of slides from individual animals with Image-Pro Plus software (Mediatech). The investigator was blinded to the experimental groups at the time of analysis. Picrosirius red staining was performed as previously described.^25,27^ In brief, quantification of interstitial staining of picrosirius red (SR) staining was performed in a blinded manner using Image-Pro software with randomly selected cortical fields. SR glomerular staining was subtracted and net SR area was normalized to net tubulointerstitial area of 400x field (Net area = Total – glomerular area - empty space). Masson Trichrome and hematoxylin eosin stains were performed on paraffin sections by standard protocols. Interstitial fibrosis and tubular atrophy (IFTA) scores was analyzed on 6 randomly selected 400x Masson Trichrome stained images. The following IFTA scores were assigned, in a blinded manner, based on the estimated percent area affected with tubular atrophy, loss of tubular structure, interstitial inflammation, and interstitial fibrosis in the field: 1 (normal); 2 (<10%), 3 (10-25%); 4 (26-50%); or 5 (>50%). Dilated tubular area was measured using Image-Pro software on 400x Masson trichrome stained images. Tubular casts and intraluminal debris area was measured using Image-Pro software on 200x PAS-stained images and normalized to net tubulointerstitial area for 6 randomly selected cortical fields. Secondary antibodies were shown to be non-reactive with tissue sections stained without the primary antibody.

### F4/80 macrophage quantification

Mice were perfused with cold normal saline and contralateral and UUO kidneys were placed on ice, digested with Liberase TL (Roche) with 1% DNase (Sigma-Aldrich), then placed at 37°C for 10 minutes, as previously described. Glomeruli were removed by passing cell suspension through a 40μm Nylon filter. Cells were stained per protocol with DAPI, PE-Cy7-anti-CD45, PE-anti-CD11b, APC-eFluor780-F4/80 from BD Sciences. Cells were blocked with mouse Fc Block (BD Biosciences). Leukocytes were identified and gated based on their positive F4/80 expression. Data was acquired on the LSR II flow cytometer (BD Biosciences) and analyzed using FlowJo software (Tree Star, Inc).

### Kidney function

Serum was analyzed for blood urea nitrogen (BUN) using the Urea Nitrogen (BUN) Reagent Set kit (Teco Diagnostics). Samples were processed according to manufacturer’s protocol. All samples were performed in triplicate.

### Immunofluoresence

For cryosectioning, excised tissue was embedded and flash-frozen in O.C.T medium (Tissue Tech) using a dry-ice slurry/2-methylbutanol mixture and cryosectioned between 8-10um. Tissue cryosections were washied with PBS and fixed with 4% PFA for 10 min. Post fixation, slides were washed three times for 5 min each with PBS followed by permeabilization using 0.2% Triton-X100 in PBS (PBT) for 10 min. Slides were then blocked (5% BSA, 2% normal goat serum in PBT) at room temperature for 1hr. Post block, tissue sections were then incubated in primary antibody overnight at 4C in blocking solution (3% BSA, 0.2% Triton-X100 in PBS). Primary antibodies used include pan-Laminin (Abcam, #ab11575), *Acta2* (Sigma, #A2547), F4/80 (Invitrogen, #MF48020), and *Cdh1* (BD Bioscience, #610181). After overnight incubation, slides were washed with PBS, and then incubated with goat ALEXA-Fluor 594- or ALEXA-Fluor 488- conjugated antibodies (Thermo Fisher Scientific) for 2h at room temperature in blocking solution. Cell nuclei were counterstained with DAPI (Molecular Probes) and mounted in 4% (w/v) propyl gallate anti-fade solution. Immunofluorescent images were obtained using an SP5 confocal microscope (Leica). *Acta2* and *Cdh1* confocal image analysis was performed as previously described.^25,27^

### Statistical analysis

All data are presented as the mean and standard error. All statistical analyses were performed using GraphPad PRISM 7.0 (GraphPad Software) and STATA 14 (StataCorp LP) software. Two-way analysis of variance (ANOVA) was performed for all parametric data including computer-assisted image analysis data for time and species. For image analysis data, the arithmetic mean of six randomly selected images of slides for each animal was used for the two-way ANOVA. Sidak’s and Tukey’s multiple comparison post-tests were utilized for time and species, respectively. Nonparametric data (IFTA and tubular injury scores) was analyzed using the Mann-Whitney U test. A *P* value <0.05 was considered statistically significant. UUO kidney weights were analyzed by linear regression.

## References

1. Duffield, J.S. et al. Host responses in tissue repair and fibrosis. Annu. Rev. Pathol. 8, 241–276 (2013).

2. Gurtner, G.C. et al. Wound repair and regeneration. Nature 453, 314–321 (2008).

3. Humphreys, B.D. Mechanisms of renal fibrosis. Annu. Rev. Physiol. 80, (2018).

4. Seifert, A.W. et al. Skin shedding and tissue regeneration in African spiny mice (*Acomys*). Nature 489, 561–565 (2012).

5. Gawriluk, T.R. et al. Comparative analysis of ear-hole closure identifies epimorphic regeneration as a discrete trait in mammals. Nat. Commun. 7, 11164 (2016).

6. Matias Santos, D. et al. Ear wound regeneration in the African spiny mouse Acomys cahirinus. Regeneration (Oxf) 3, 52–61 (2016).

7. Duffield, J.S. Cellular and molecular mechanisms in kidney fibrosis. J. Clin. Invest. 124, 2299–2306 (2014).

8. Tapmeier, T.T. et al. Reimplantation of the ureter after unilateral ureteral obstruction provides a model that allows functional evaluation. Kidney Int. 73, 885–889 (2008).

9. Liu, J. et al. Cell-specific translational profiling in acute kidney injury. J. Clin. Invest. 124, 1242–1254 (2014).

10. Lin, S.L. et al. Bone marrow Ly6Chigh monocytes are selectively recruited to injured kidney and differentiate into functionally distinct populations. J. Immunol. 183, 6733–6743 (2009).

11. Liu, W. et al. Dragon (repulsive guidance moledule GGMb) inhibits E-cadherin expression and induces apoptosis in renal tubular epithelial cells. J. Biol. Chem. 288, 31528–31539 (2013).

12. Chaabane, W. et al. Renal functional decline and glomerulotubular injury are arrested but not restored by release of unilateral ureteral obstruction (UUO). Am. J. Physiol. Renal Physiol. 304, F432–F439 (2013).

13. Zheng, G. et al. Alpha3 integrin of cell-cell contact mediates kidney fibrosis by integrin-linked kinase in proximal tubular E-cadherin-deficient mice. Am. J. Pathol. 186,1847–1860 (2016).

14. Leferovich, J.M. et al. Heart regeneration in adult MRL mice. Proc. Natl. Acad. Sci. USA 98, 9830–9835 (2001).

15. Robey, T.E., Murry, C.E. Absence of regeneration in the MRL/MpJ mouse heart following infarction or cryoinjury. Cardiovasc. Pathol. 17, 6–13 (2008).

16. Oh, Y.S. et al. Scar formation after ischemic myocardial injury in MRL mice. Cardiovasc. Pathol. 13, 203–206 (2004).

17. Iwata, T. et al. Aberrant macrophages mediate defective kidney repair that triggers nephritis in lupus-susceptible mice. J. Immunol. 188, 4568–4580 (2012).

18. Chawla, L.S., et al. Acute kidney injury and chronic kidney disease as interconnected syndromes. New Engl. J. Med. 371, 58–66 (2014).

19. Ferenbach DA, Bonventre JV. Mechanisms of maladaptive repair after AKI leading to accelerated kidney ageing and CKD. Nat. Rev. Nephrol. 11, 264–276 (2015).

20. Liu, J. et al. Molecular characterization of the transition from acute to chronic kidney injury following ischemia/reperfusion. JCI Insight 2, e94716 (2017).

21. Poss, K.D. Advances in understanding tissue regenerative capacity and mechanisms in animals. Nat. Rev. Genet. 11, 710–722 (2010).

22. McCampbell, K.K., Wingert, R.A. New tides: using zebrafish to study renal regeneration. Transl. Res. 163, 109–122 (2014).

23. Kragl, M. et al. Cells keep a memory of their tissue origin during axolotl limb regeneration. Nature. 460, 60–65 (2009).

24. Kikuchi, K. et al. Primary contribution to zebrafish heart regeneration by gata4(+) cardiomyocytes. Nature. 464, 601–605 (2010).

25. Pennathur, S. et al. The macrophage phagocytic receptor CD36 fibrogenic pathways on removal of apoptotic cells during chronic kidney injury. Am J Pathol 185, 2232–2245 (2015).

26. Okamura, DM. et al. Cysteamine modulates oxidative stress and blocks myofibroblast activity in CKD. J Am Soc Nephrol 25, 43–54 (2014).

27. Okamura, DM. et al. CD36 regulates oxidative stress and inflammation in hypercholesterolemic CKD. J Am Soc Nephrol 20, 495–505 (2009).

